# Climate regulation of fruit-set, orchard synchrony, and future production in apple agroecosystems across the Republic of Korea

**DOI:** 10.64898/2026.08.18.745556

**Authors:** Arne Buechling, Jung Gun Cho, Jakob Pavlin, Forzia Ibrahim, Patrick H. Martin

## Abstract

A better understanding factors regulating agricultural production is a research priority, given the pace of environmental change and global human population growth. Fruit orchards may be particularly vulnerable to climate change owing to the strong temperature-dependency of reproduction in temperate trees. In this study, we explored climate influences on the reproductive dynamics of apple (*Malus domestica*), one of the most widely-grown, economically-important fruit crops worldwide. Observations of individual-tree fruit-set, a key indicator of final yield, were acquired for three apple cultivars in an eight-year census of ∼44,900 trees in >5,500 orchards spanning wide climate gradients across the Republic of Korea. With maximum-likelihood models, we quantified temporal and climate-driven patterns in fruit-set, investigated evidence for spatially-synchronized production, and conducted simulations of orchard vulnerability to climate change. We found annual fruit-set oscillated around modal levels and was synchronized between orchards within 25 km. Higher temperatures had contrasting influences, reducing fruit-set during the cold-season (consistent with climate-induced phenological shifts), while increasing fruit-set during the season of bud initiation (prior spring). Simulations predict that higher future temperatures during bud initiation increase average fruit-set, despite the constraints of cold-season warming, but that such enhancements level-off by late-century and are accompanied by heightened volatility. Decelerating fruit-set and greater instability in production have implications for future societal needs, as robust supply systems depend as much on consistent production as on high average output. Our analyses also imply that future climate patterns promoting synchrony in fruit-set among orchards may accentuate the negative consequences of fluctuating production.

## Introduction

In an era of rapid global change, assessing the capacity of agricultural systems to support the food demands of a growing world population is a research priority (Tilman, 2022). Increasing frequencies of severe drought and anomalous temperatures, considered signatures of anthropogenic climate change (Horton et al., 2016), are expected to threaten agricultural output and compromise global food supplies (Springmann et al., 2016; Heino et al., 2023; Novick et al., 2024; Bellvert et al., 2025). While farm productivity has markedly improved in the past half-century (Godfray et al., 2010), recent evidence points to a slowing of yield growth in some important food crops (Lobell et al., 2011; Ray et al., 2012; Kucharik et al., 2020; Gleiser et al., 2021). Decelerating yields are particularly evident for high-value perennial crops, such as orchard trees that are cultivated for seed and fruit production (Aizen et al., 2023), and the negative effects of a slowing of crop growth are compounded by instances of severe crop failures as recent extreme heat events have triggered outright reductions in orchard yields in some regions (Parker et al., 2020; Preston et al., 2024). These trends are of societal concern, as cultivated fruit is a key component of human diet and nutrition, providing essential vitamins, minerals, and fiber (Bondonno et al., 2017; İkinci, 2025). The apple (*Malus domestica* Borkh.), in particular, is globally important in terms of production, consumption, agriculture employment, and international trade (Cornille et al., 2014; González Noguer et al., 2023; Chen et al., 2025). However, as a majority of research concerning climate change impacts on food production has focused on cereal crops, considerable knowledge gaps remain regarding the long-term outcomes for perennial crops such as apple (Parker et al., 2020; Meza et al., 2023; Pickson et al., 2026).

We suggest that more robust forecasts of the climate-change consequences for the viability and diversity of global agricultural supply will be supported by an improved understanding of the climate relations of food plants beyond cereals (Wheeler and Von Braun, 2013). The evidence to date indicates that fruiting in tree crops of temperate regions is influenced by complex interactions between climate and plant physiology that span multiple seasons within their near year-long reproductive cycles (Schauberger et al., 2016). For example, a large body of work has investigated the temperature-sensitivity of the phase of bud dormancy, during which metabolic processes are inhibited to facilitate cold resistance (Campoy et al., 2011). Low temperatures (chilling) in mid-winter combined with progressive warming (forcing) in spring are understood to promote a gradual reactivation of developmental and vascular functions within buds (Savage and Chuine, 2021). For many cultivars, the magnitude, duration, and interrelationships of their chilling and forcing requirements have not been reliably established (Laub et al., 2014; Chuine et al., 2016; Wang et al., 2020; Delgado et al., 2021). In general however, global warming is expected to reduce chilling but enhance forcing, ostensibly explaining widespread observations for an advancement of budbreak in many species (Menzel et al., 2006; Legave et al., 2013; Piao et al., 2019; Ettinger et al., 2020; Cho et al., 2021). In the context of fruit-tree productivity, the focus of this study, cold-season warming trends that disrupt dormancy have been associated with erratic flowering and significant reductions in fruit-set (Faust et al., 1997; Atkinson et al., 2013; Campoy et al., 2019; Roussos, 2024).

Reproductive output in plants also depends on the availability of resources, the acquisition of which is similarly mediated by climate (Roussos, 2024). Temperature, for example, both directly and indirectly affects assimilation processes and the magnitude of carbohydrate reserves necessary for tissue growth. Warming enhances enzyme function and photosynthetic rates up to physiological optimum levels, while excessive heat can increase respiration demands, degrade proteins, and limit net carbon gains (Saxe et al., 2001; Dusenge et al., 2019). High temperatures also accelerate soil drying and amplify vapor pressure deficits, which can elevate tension in the water-conducting vasculature (xylem) of trees, potentially impairing transport functions (Tyree and Sperry, 1989; Grossiord et al., 2020). Compromised hydraulic capacity can trigger stomatal closure in leaves, reducing gas exchange, and therefore, photosynthetic capacity (Ryan and Yoder, 1997). Further, as climate tends to be correlated over large areas, climate variation may impose similar fluctuating constraints on the physiological performance and resources of widely-dispersed trees, potentially leading to a synchronization of fruiting among disjunct orchards (Crone and Rapp, 2014). The degree and drivers of spatially correlated reproductive effort have been widely investigated for wind-pollinated plants in natural ecosystems (masting research), but comparatively less is known about the potential for synchronized processes in orchards and for insect-pollinated taxa (Garcia et al., 2021).

Additionally, variation in crop size may be determined not only by phenology and resources, as influenced by climate, but also by fundamental tradeoffs in the allocation of resources to different biological processes (Obeso, 2002). An inherently finite supply of carbon and nutrients necessitates a balancing of investments in reproduction versus functions related to survival and growth (Bazzaz et al., 1987). Models have shown that allocation to growth, defense and basic tissue maintenance is prioritized in perennial plants, whereas disbursements to fruiting depend on the availability of surplus resources that fluctuate across years (Isagi et al., 1997). The influence of climate coupled with intrinsic allocation tradeoffs may therefore operate in concert to further enforce a pattern of oscillating reproductive output. Crop production that fluctuates annually and is potentially synchronized among orchards has implications for societal food security, which is underpinned by an adequate mean supply rate from farms, as well as a low variance in supply across years (Schmidhuber and Tubiello, 2007; Garibaldi et al., 2011).

In this study, we investigate the dynamics of fruit-set in apple, assess evidence of covarying levels of production among disjunct orchards, and quantify relationships with climate. We use annual-resolution records of per capita fruit-set for three common cultivars grown in orchards spanning extensive geographical and climate gradients across the Republic of Korea (Appendix A1). Fruit-set integrates tree carbon balance, phenology, blossom production, pollination processes, initial abortion of fertilized ovaries, and the effects of current and prior climate, and is thus a robust indicator of reproductive dynamics. Fruit-set is also strongly correlated with total crop yield (Brown, 1952; Forshey and Elfving, 1977; Brain and Landsberg, 1981; Dennis, 1981; Atkinson et al., 2013; Monselise, 2018; González-Martínez et al., 2026). Our goal was to model fruit-set capacity and not phenological processes such as budbreak *per se*, but our approach implicitly integrates the influence and consequences of shifting patterns of phenology observed both globally and in the Republic of Korea (Cho et al., 2021). To understand production-climate relations, we adopt a nonparametric analysis framework often used to model tree demographic processes (reproduction, growth, survival) in natural forested systems (e.g., Canham and Uriarte, 2006; Gómez-Aparicio et al., 2008, 2011; Buechling et al., 2016; Korolyova et al., 2022; Ibrahim et al., 2025). Within this framework, we formulate nonlinear models that quantify the magnitude and shape of climate effects associated with multiple stages of the reproductive cycle of apple. We expect, based on findings from the phenology literature, that anomalous warming during the period of winter dormancy compromises fruit-set capacity. We also posit that fruit-set production is tempered by climate conditions in seasons preceding bud dormancy through effects on plant carbon balance. We additionally conduct spatially-explicit simulations at a nationwide scale to ask whether future warming will cause production changes, if thresholds exist that reserve of any gains, if climate change destabilizes production, and if outcomes vary geographically.

## Methods

### Study region and production data

The climate of the study region is temperate and influenced by the East Asian monsoonal circulation system. The growing season spans months from March to August and typically receives a majority (∼900 mm) of total annual precipitation (∼1200 mm). Mean growing season temperatures range from 11.0-12.5°C.

We acquired records of fruit-set from a private company, NongHyup Property and Casualty Insurance. The company initiated a field-based monitoring program in 2005 to document annual variation in production in apple orchards in terms of fruit-set quantities. A stratified random design was used to select a set of orchards for sampling that spanned the full range of environments suitable for apple cultivation in the Republic of Korea. Field surveys were conducted from late June to early July of each year. Within orchards, 10–25 trees were randomly selected for sampling. Tree ages were identified from records of planting dates (Table 1).

**Table 1.**
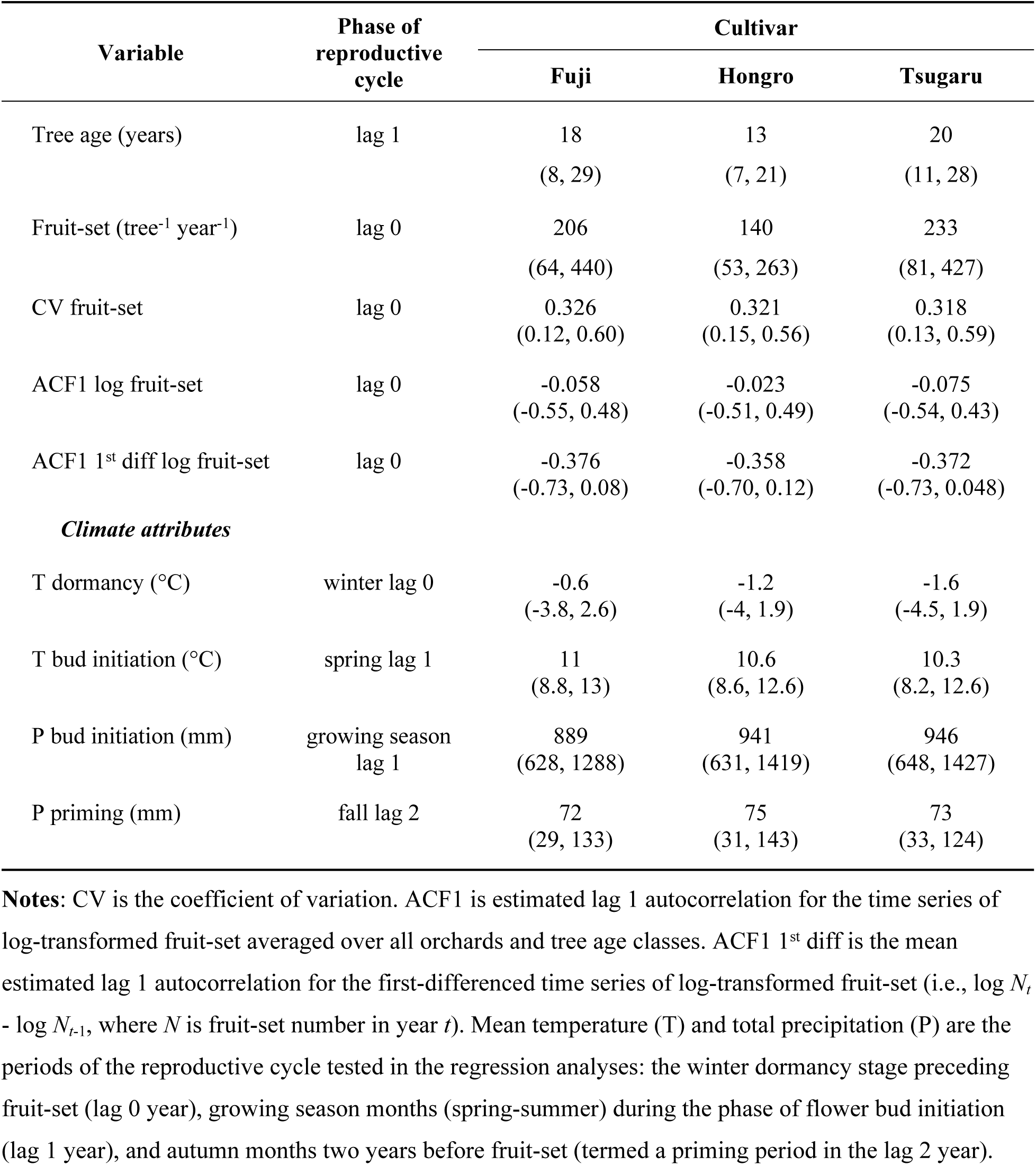
Attributes of model-fitting datasets, including observed fruit-set per tree and climate variables summarized over the analysis window (2007-2014). Mean values are shown with 5^th^ and 95^th^ percentiles, representing orchard-level variability (accounting for age class) in brackets. Lag 0 is the fruit-set year, lag 1 is the bud initiation year, and lag 2 is two years prior to fruiting.

We analyzed patterns of annual fruit-set for three cultivars (Fuji, Hongro, Tsugaru) over an eight-year period (2007-2014). Orchard records with missing data were excluded. The individual tree data were averaged to produce separate chronologies of annual, per capita fruit-set for each cultivar and age class present in each orchard. Final sample sizes of orchards were 3990, 884, and 655 for Fuji, Hongro, and Tsugaru, respectively (Fig. 1).

**Figure 1.**
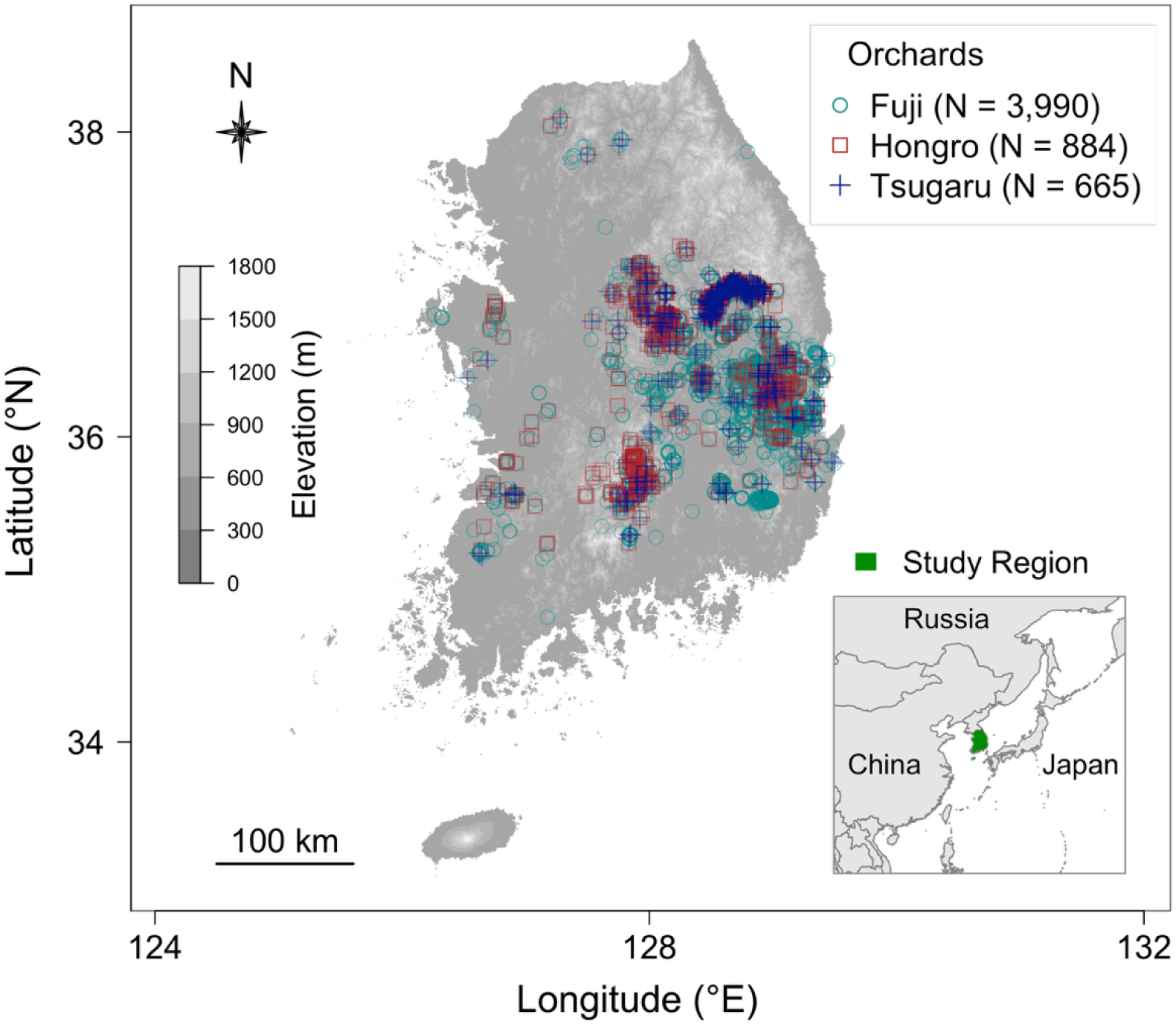
Location and sample size (N) of surveyed orchards in the Republic of Korea for three apple cultivars plotted against a 30 m digital elevation model.

Standard descriptive and nonparametric statistics were computed to understand temporal and spatial patterns in fruit-set. We generated histograms and conducted Anderson-Darling tests to determine whether data were normally distributed. Following approaches from the forest ecology literature (Koenig, 2021), we computed the coefficient of variation and autocorrelation functions (ACF) to assess the degree and direction of temporal variation in fruit-set abundances. We calculated ACFs with a lag of one year for the time series of log-transformed fruit-set, as well as the first-differenced chronologies of log-transformed fruit-set. First-differencing is a detrending step that focuses the analysis on directional changes in fruit-set, rather than variation in absolute quantities (Liebhold et al., 2004). To investigate the extent that annual reproductive output covaried or was synchronized among distant orchards, we fit nonparametric spline correlograms (Bjørnstad and Falck, 2001) to the spatially-explicit chronologies of fruit-set counts.

### Climate data

We used geostatistical techniques to produce continuous, gridded datasets of temperature and precipitation for the study area. We acquired instrumental climate records from 88 widely-distributed weather stations from the Korea Meteorological Administration. Temperature was estimated using inverse distance weighting (IDW). Temperature values for all individual raster cells were estimated by weighting and then averaging instrumental records from the nearest adjacent weather stations. The weighting scheme was dependent on and inversely proportional to the Euclidean distance between a given grid cell and the closest weather station locations (Shepard, 1968). Grid-cell estimates were adjusted for elevation-dependent pressure gradients using a fixed lapse rate correction (6.5°C km^−1^) (Tercek et al., 2021). We used a 30 m digital elevation model (Tachikawa et al., 2011) to determine grid-cell elevations.

For the spatial interpolation of precipitation, we used ordinary kriging. Kriging has previously been found to provide more accurate estimates of precipitation than IDW (Goovaerts, 2000; Mair and Fares, 2010). Similar to IDW, kriging computes a weighted average estimate based on distances to station observations, but weights are derived from measures of dissimilarity or spatial autocorrelation between the station data. We estimated patterns of spatial autocorrelation by constructing a semi-variogram fit with a continuous spherical function.

The resulting interpolated climate grids had a 300 m spatial resolution. We then extracted climate values for each orchard location using bilinear interpolation. Monthly data were aggregated to generate annual and seasonal estimates of mean temperature and total precipitation for each year of the analysis period. Interpolation analyses were performed in ArcGIS (10.3, ESRI, Redlands, California).

### Regression analyses

We formulated sets of cultivar-specific regression models to evaluate the effects of climate on fruit-set. Apple trees flower and set fruit annually, but full floral development (from bud primordia initiation to anthesis) spans a time period of ∼9 months that includes a stage of cold-season dormancy (Pratt, 1988). We assumed that the climate conditions over the seasons that span the floral developmental period set the fundamental constraints on fruit-set potential. Our fruit-set models therefore incorporated climate factors representing multiple, fixed seasonal intervals that were selected *a priori*. The models were formulated to assess evidence for the following specific hypotheses: (1) warming during the cold-season (December-February) reduces fruit-set by disrupting dormancy processes; (2) increases in prior-year spring (March-May) temperatures and growing-season (March-August) precipitation enhances fruit-set up to threshold levels set by cultivar-specific photosynthetic and physiological optima; and (3) fall (September-November) climate two years before fruit-set ‘primes’ a tree’s future fruit-set capacity by influencing the levels of carbohydrate reserves available to support metabolic activity in succeeding seasons.

We assumed that local management factors are important, but we lacked orchard-specific information regarding cultivation practices, a common limitation associated with large-scale datasets of agricultural production (Gleiser et al., 2021). Therefore, we generated a model covariate to serve as a proxy (*PotFruit*, Eqn. 1) for management and other orchard-scale factors that may modify fruit-set capacity; specifically, to account for factors not explained by the fixed effects of our models, we estimated unique levels of maximum fruit-set potential for groups of orchards classified according to congruent patterns of observed fruit-set. We used agglomerative hierarchical clustering with a complete linkage function for the classification. Four groups were identified that minimized within-group and maximized between-group variability.

Our modeling approach is grounded in likelihood theory (Edwards, 1992). We used an iterative optimization algorithm (see section 2.4) to estimate the maximum-likelihood values of the unknown parameters of a model. Separate sets of nonlinear models were fit for each cultivar. Models were multiplicative in form:

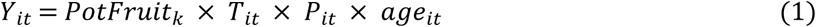

where *Y_it_* is expected per capita fruit-set in orchard *i* in year *t*. *PotFruit* is estimated maximum fruit-set potential, *T* is mean temperature for a given stage of the reproductive cycle, *P* is total precipitation for a particular stage, and *age* is tree age in year *t*. We estimated unique levels of *PotFruit_k_* for *K*=4 orchard groups. In effect, this regression framework produces multi-level models that account for orchard-specific variation in fruit-set capacity. Covariates (functional forms below) are scalar quantities (values range from 0-1) that proportionally decrease maximum fruit-set potential (*PotFruit*). Scalar forms mitigate collinearity and parameter trade-off issues, and promote model convergence (Canham and Murphy, 2016).

We used nonlinear functions to estimate relationships between fruit-set and explanatory variables. We tested lognormal and Gaussian functions, each capable of fitting monotonically increasing, decreasing, or unimodal distributions. Seasonal temperature effects were modeled using symmetrical Gaussian functions:

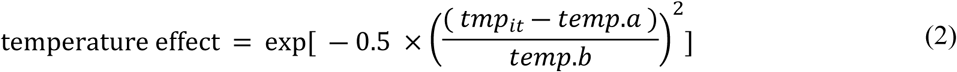

where *tmp_it_*is observed mean temperature (°C) for a specific season in orchard *i* in year *t*, and *temp.a* and *temp.b* are the mode (location) and variance of the Gaussian, respectively.

Seasonal precipitation effects were fit with asymmetrical lognormal functions:

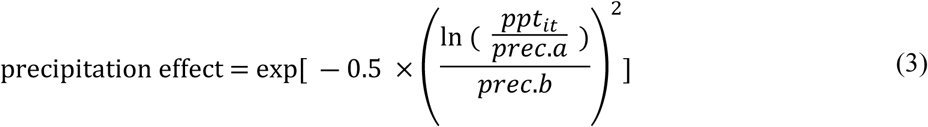

where *ppt_it_* is observed total precipitation (mm) in a given season for orchard *i* in year *t*, and *prec.a* and *prec.b* describe the mode and breadth of the lognormal. We also estimated the effect of tree age using a lognormal function of the same form as Eqn. 3.

### Model specification

We used simulated annealing (Goffe et al., 1994), a global optimization algorithm based on the Metropolis criterion, to solve for the maximum-likelihood estimates of model parameters (Chuine et al., 1998). To account for heteroscedasticity, likelihood was computed using a modified normal probability density function where variance was a linear function of the mean. We quantified parameter uncertainty using asymptotic support intervals, defined as the range in parameter values corresponding to a two-unit change in likelihood (Edwards, 1992). Goodness of model fit was assessed based on visual examination of observed versus predicted values, as well as measures of bias, mean absolute error, and *R^2^*. Bias was determined from the slope of the regression of observed against predicted values (see Table 2 for details). In this study, *R^2^* was computed from residual sums of squares (*R^2^*=1-sum of squares error/sum of squares total), representing the fraction of the total sum of squares explained by a model (Cameron and Windmeijer, 1997). *R^2^* is bounded by 1, when observations precisely match expected values given the model, and declines as the average deviation from expected values increases (Buechling et al., 2026b). Negative *R^2^* values indicate that deviations from expected values are on average greater than the deviations from the simple mean of the observations, indicating that model predictions are no better than simply using the mean of the observations (Cook et al., 1987; Clark et al., 1998). The degree of support for a given model was assessed using Akaike information criterion (AIC) (Burnham and Anderson, 2002). AIC values were adjusted (AICc) to account for model complexity by integrating a bias correction term (Johnson and Omland 2004). Analyses were conducted in R (R Core Team, 2025). Models were fit using the likelihood package (Murphy, 2026).

**Table 2.**
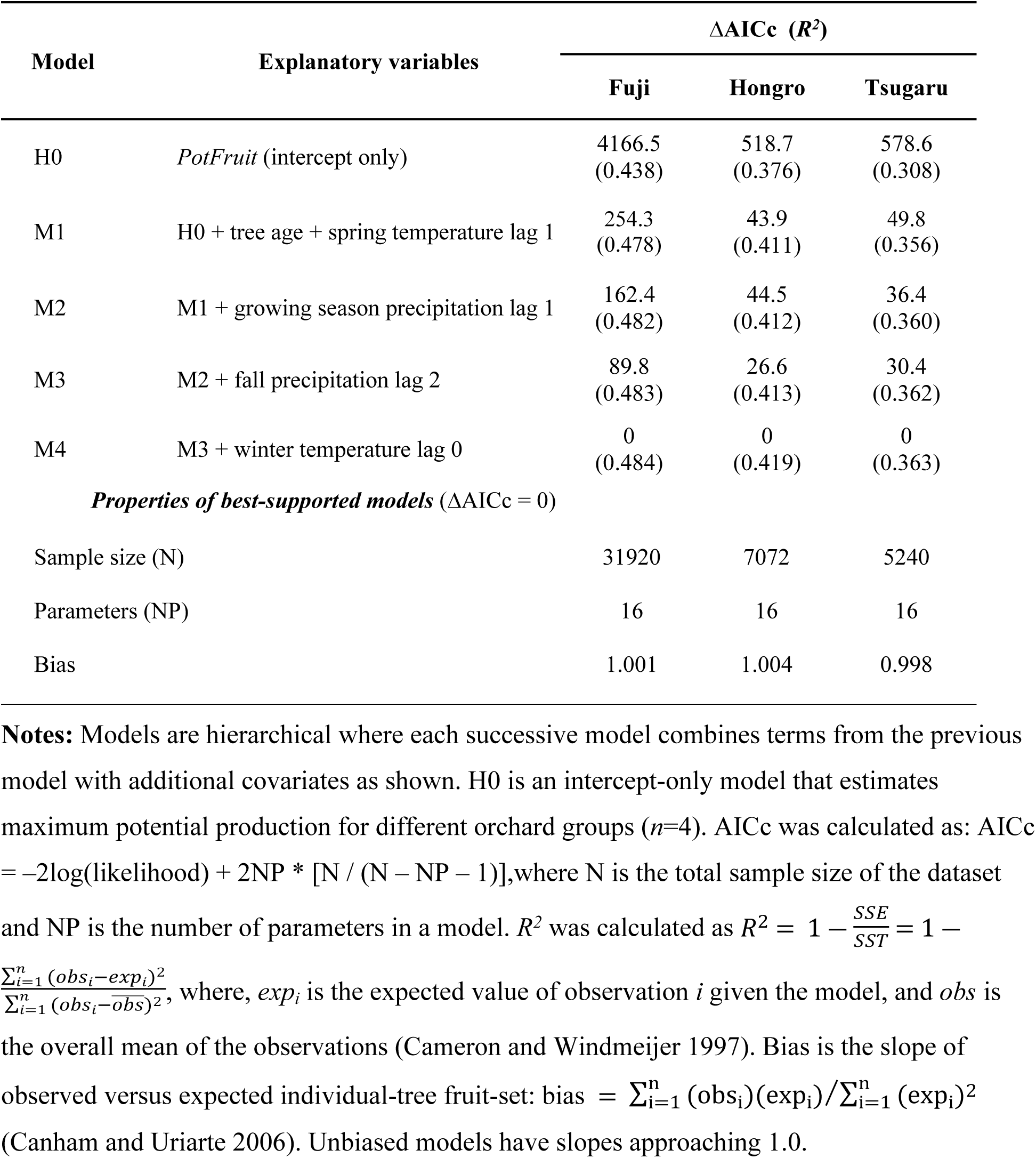
Fruit-set production model results for each cultivar. Models were compared per AIC values corrected for model complexity (AICc). ΔAICc is the difference between AICc of a given model and the minimum AICc of the model set. *R^2^* values are shown in brackets. Lag 0 is the fruit-set year, lag 1 is the bud initiation year, and lag 2 is two years prior to fruiting.

### Simulations

We used the best-supported (per AICc) fecundity models for each cultivar to predict fruit-set responses to projected future climate conditions. Simulation models were fit with temperature and precipitation datasets generated from regional-scale climate models (CM) based on different scenarios of socio-economic development and carbon emissions (van Vuuren et al., 2011). These scenarios, termed relative concentration pathways (RCP), describe factors leading to particular ranges of radiative forcing values expected by year 2100. We used CM output data associated with a moderate climate change scenario that leads to atmospheric CO_2_ concentrations of ∼650 ppm by 2100 (RCP4.5), and an extreme development trajectory that results in CO_2_ concentrations of ∼1370 ppm (RCP8.5). Climate projection data were acquired from the Korea Meteorological Administration for the years 2023-2100 and integrated output from an ensemble average of five independent regional CMs (Suh et al., 2012). The climate datasets were continuous rasters with a monthly resolution. In ArcGIS, we downscaled the rasters from 1.0 km to 300 m to match the resolution of the interpolated climate grids.

Simulations estimated spatial-temporal patterns of fruit-set for a hypothetical individual tree over an ∼80-year period ending in 2100. We integrated climate variables for multiple phases of the reproductive cycle (priming, bud initiation, and winter dormancy; Appendix A2). Simulations were conducted to predict fruit-set for each pixel within the extent of mainland Korea. We compared mean levels of predicted fruit-set for three 30-year periods: near future (2031-2060), far future (2071-2100), and a reference interval (1985-2014). Predicted fruit-set in the reference interval was derived from regression models fitted with interpolated climate values, rather than the CM data.

To explore temporal patterns, we derived, following Gleiser et al. (2021), estimates of the mean annual growth rate of fruit-set over successive years (*Δfruitset*) and the corresponding standard deviation of growth (*SDΔfruitset*). Mean growth rate was computed from the first-differenced time series of fruit-set for a given orchard:

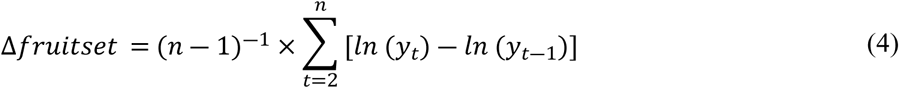

where *Δfruitset* is the mean difference of log-transformed annual fruit-set (*y*) between successive years (*t*-1 and *t*), and *n* is the number of observations in a time series for a given cultivar, orchard, and age class. Standard deviation of Δfruitset was then calculated as:

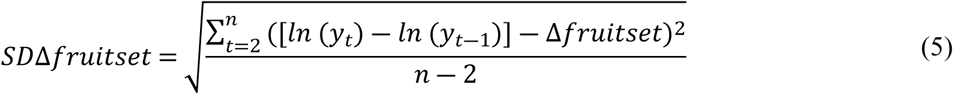

where *SDΔfruitset* is a linearly detrended measure of interannual variability in log-transformed fruit-set (Gleiser et al., 2021).

## Results

We computed various metrics to characterize the temporal and spatial dynamics of production. Histograms of fruit-set (Appendix A3, Fig. A3.1) indicate that distributions were unimodal and marginally right-skewed. Accordingly, the arithmetic means of the data were consistently greater than the corresponding medians and the Anderson–Darling tests were significant (p<0.001) for cultivars, indicating a departure from normality.

Observed levels of annual fruit-set varied considerably between years, ranging from 53-440 fruit per tree depending upon cultivar (Table 1). Within a given cultivar and orchard group, the magnitude of fruit-set in successive years deviated by up to ∼30% relative to overall mean levels (Fig. 2). The coefficient of variation (CV) of annual untransformed fruit-set, averaged over all orchards, was ∼0.32 for all cultivars (Table 1). The frequency distribution of orchard-scale CVs was relatively broad and marginally right-skewed with maximum values of ∼0.9 (Appendix A3, Fig. A3.2).

**Figure 2.**
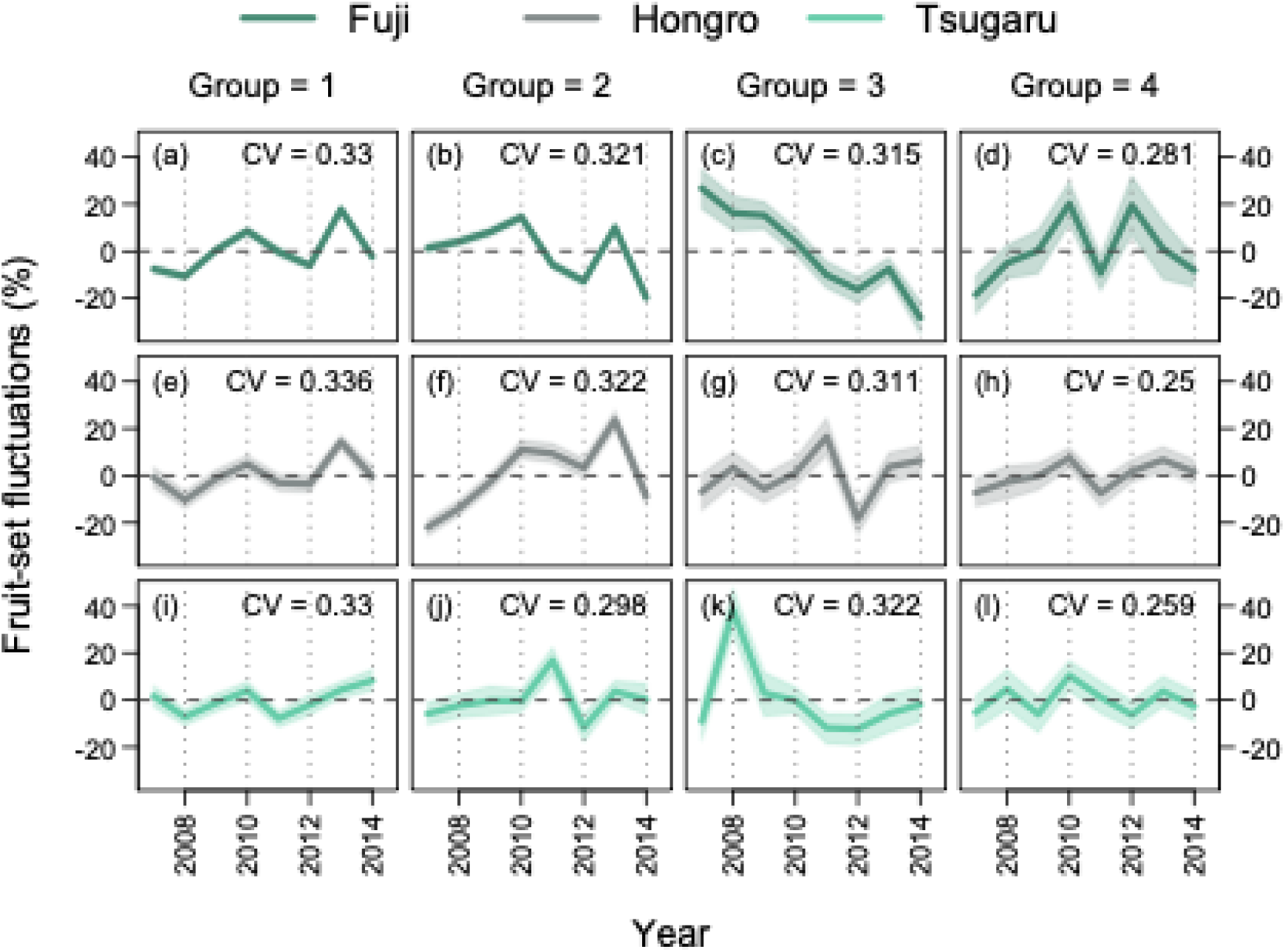
Time series of observed fluctuations in annual per capita fruit-set by cultivar and management group. Shown is the annual percent deviation of fruit-set above or below the overall mean level of a given cultivar and orchard group. The dashed zero reference line is the mean level of untransformed fruit-set. CV is the coefficient of variation. Shaded regions are bootstrapped 95% confidence limits.

Negligible levels of temporal serial autocorrelation were observed in the time series of log-transformed fruit-set (Table 1). In contrast, stronger (though generally non-significant) levels of negative temporal autocorrelation at one-year lags were revealed in the first-differenced chronologies (Appendix A4). Specifically, mean orchard-scale ACF values for one-year lags were -0.38, -0.36, and -0.37 for Fuji, Hongro and Tsugaru, respectively.

Spline correlograms fit to log-transformed time series of fruit-set revealed significant spatial correlation (synchronization) among disjunct farms. Significant synchrony was detected among orchards within 25 km for Fuji and Hongro, and ∼15 km for Tsugaru (Appendix A5).

### Regression models

Consistent with expectations, orchard group, assumed to at least partially account for management factors, was a main determinant of per capita fruit-set based on *R^2^* values (model H0, Table 2). After controlling for orchard group, our analyses indicate that tree age and climate additionally mediated fruit-set. The best-supported models provide modest but reasonable levels of agreement with observational datasets, particularly given the broad environmental and climate gradients encompassed by the surveyed orchards. For Fuji, Hongro, and Tsugaru, respectively, *R^2^* values are 0.484, 0.419, and 0.363 (Table 2) and mean absolute errors are 63, 38, and 65 fruit tree^−1^ (Appendix A6, Fig. A6.1). Model predictions are unbiased (slope of observed vs. predicted ∼1.0) and residuals randomly distributed (Appendix A6, Fig. A6.2). Model parameter values and their support intervals are in Appendix A7.

Age effects on fruit-set were unimodal in shape for both Hongro and Tsugaru (Appendix A8). Production was maximized in trees approaching ∼22 and 25 years of age, controlling for other factors. Fruit-set in Fuji increased monotonically with age without reaching a threshold maximum, within the range of the model-fitting datasets.

Climate factors integrated over multiple seasons across three years were important for fruit-set capacity (model M4, Table 2). Climate during the year of bud initiation (Fig. 3) was the most important correlate of production; specifically, the combined effects of temperature in spring and total precipitation for the growing season regulated levels of fruit-set in the following year. Warmer-than-average conditions overlapping with the phase of bud initiation enhanced reproductive effort: temperature increases of 1.0-1.5 standard deviation (SD) above observed mean levels for a given cultivar increased fruit-set by up to two SDs (Fig. 4). Controlling for temperature, responses to growing-season precipitation were approximately unimodal for Fuji and Tsugaru (Fig. 3b). Subsequent fruit-set increased with moisture up to a threshold level (∼900 mm), but were depressed by precipitation beyond this limit.

**Figure 3.**
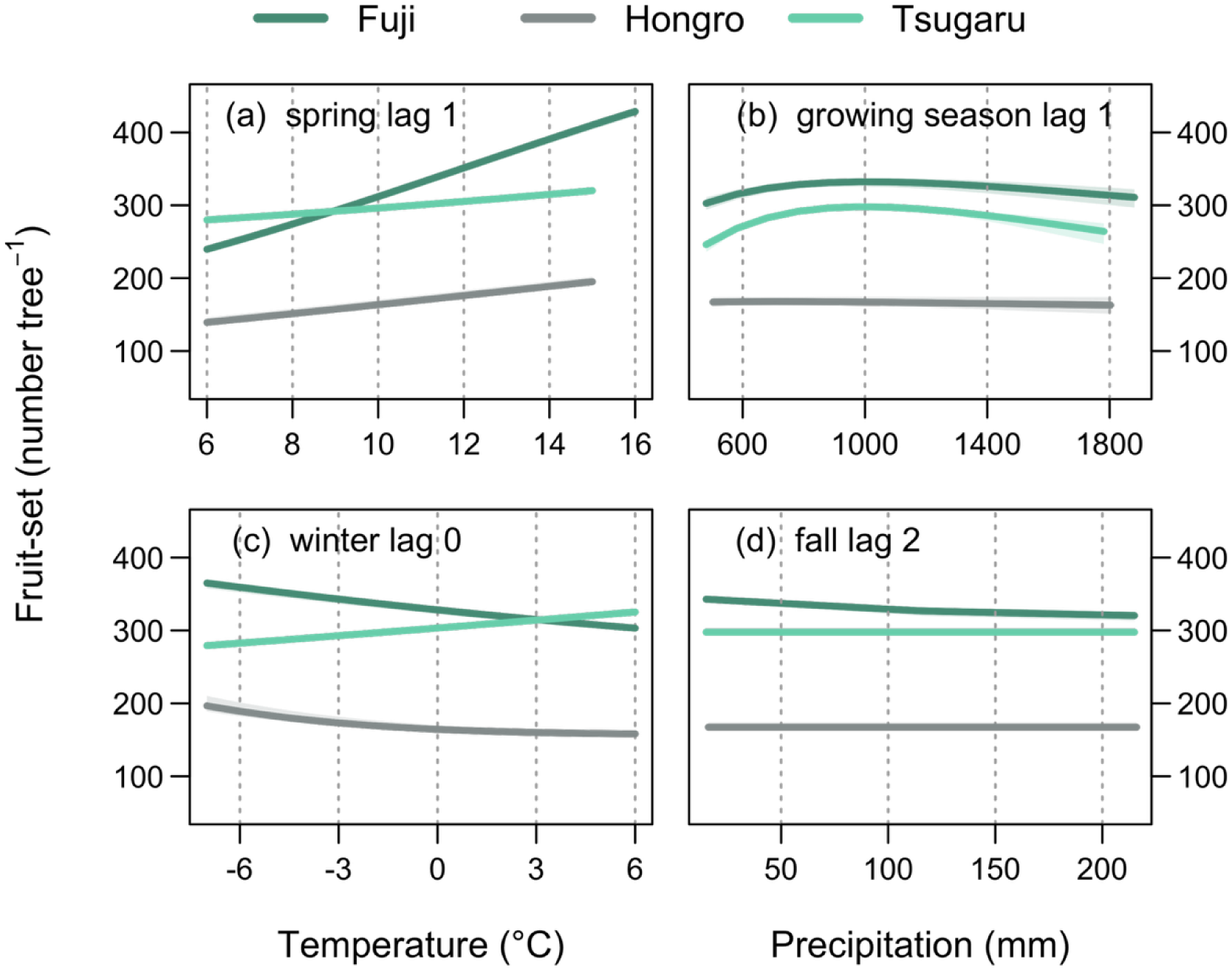
Estimated effects of climate on per capita fruit-set for three cultivars. The best-supported models incorporated climate effects for four seasons over a three-year period: (a) mean temperature for the months March-May of the bud initiation year (spring lag 1 year), (b) total precipitation from March-August during the bud initiation year (growing season lag 1 year), (c) mean temperature for the months December-February preceding flowering and fruit-set (winter lag 0 year), and (d) total precipitation for the months September-November two years prior to fruit-set (fall lag 2). Each panel shows the marginal effect of a given covariate, setting other variables in the model at their mean values. Narrow shaded regions are 2-unit asymptotic support intervals. Support limits are values above and below the maximum likelihood estimate that cause the likelihood to drop by 2 units.

**Figure 4.**
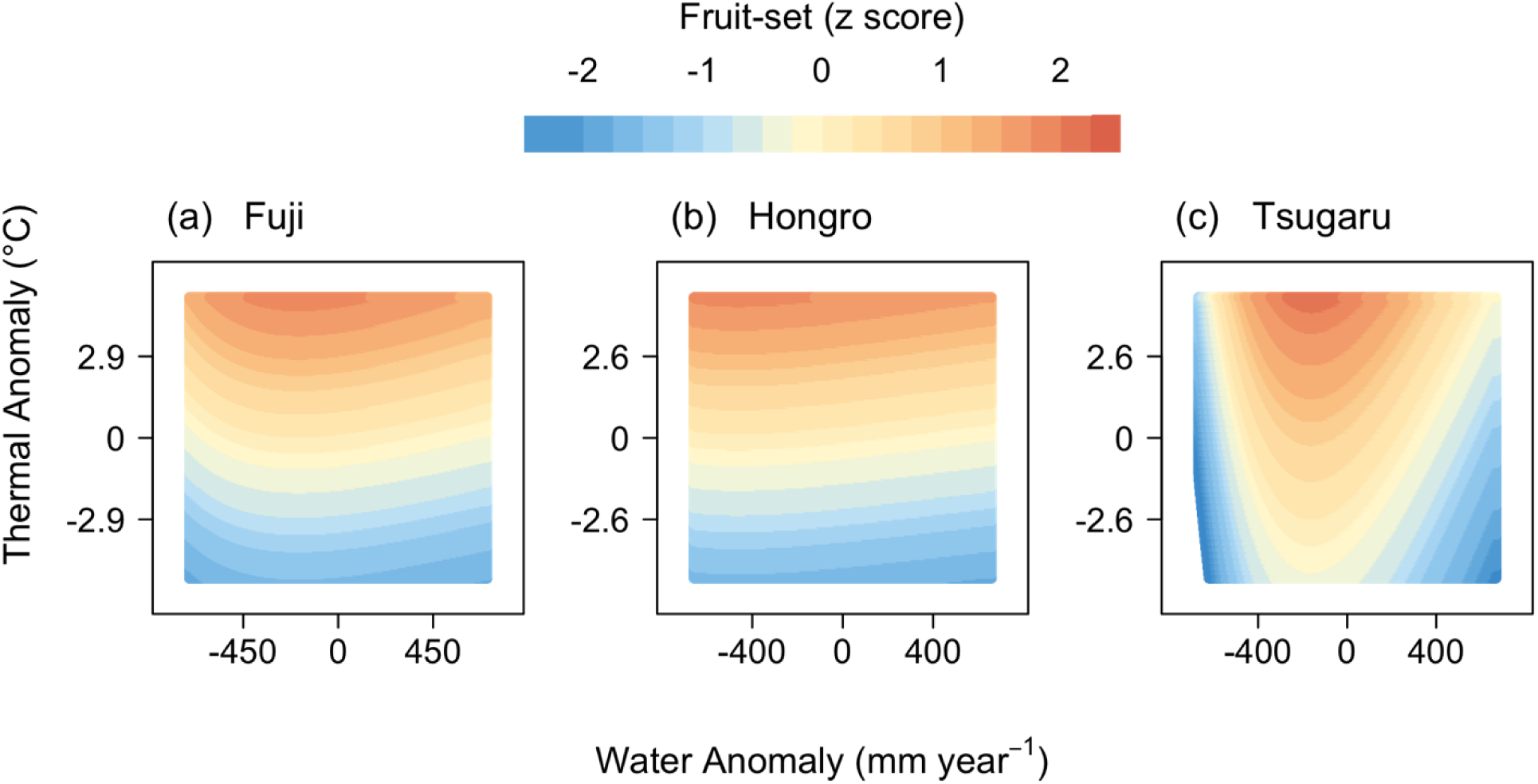
Predicted annual fruit-set in an individual tree in response to the joint, nonlinear effects of spring temperature (March-May) and growing-season precipitation (March-August) during the bud initiation year (year prior to fruit-set). Both thermal and water supply levels are shown as anomalies or departures from the corresponding mean levels for a cultivar. Climate means are scaled to zero. Climate levels that are 1.0 standard deviation above and below a mean value are labeled on each axis. Expected annual fruit-set quantities are shown as z scores (standard deviations above or below mean levels for a given cultivar).

Cultivars diverged in their sensitivity to cold-season temperatures within the bud dormancy phase preceding fruit-set (Fig. 3c). Fruit-set decreased monotonically with winter warming in Fuji and Hongro, but increased in Tsugaru.

Finally, fall precipitation two years prior to fruit-set was an important predictor of subsequent production, per AIC-based selection criteria (Table 2). However, the magnitude of this effect was negligible, as responses were generally flat (Fig. 3d).

### Simulations

Simulations derived from the best-supported models predict generally positive trends and sizeable increases in average per capita fruit-set for all cultivars under both the intermediate (RCP4.5) and high-end climate change scenario (RCP8.5). Fruit-set is forecast to increase continuously over the full simulation period (2023-2100) under RCP8.5, but plateau ∼2070 under RCP4.5 (Fig. 5). Correspondingly, larger proportional increases are forecast with RCP8.5, relative to RCP4.5, for the last three decades of the simulation period (far future, 2071-2100, Appendix A9). Specifically, under RCP8.5, annual average production in the far future period is predicted to increase by 15.5%, 8%, and 9% in Fuji, Hongro, and Tsugaru, respectively, above reference period (1984-2014) levels. Under RCP4.5, predicted changes for the far future are lower but still sizeable, increasing by 8%, 4% and 4% in Fuji, Hongro, and Tsugaru, respectively, relative to reference levels. Simulated fruit-set levels for the near future period (2031-2060) are similar under either climate change scenario, increasing by ∼4.5% in Fuji and by <3% for both Hongro and Tsugaru.

**Figure 5.**
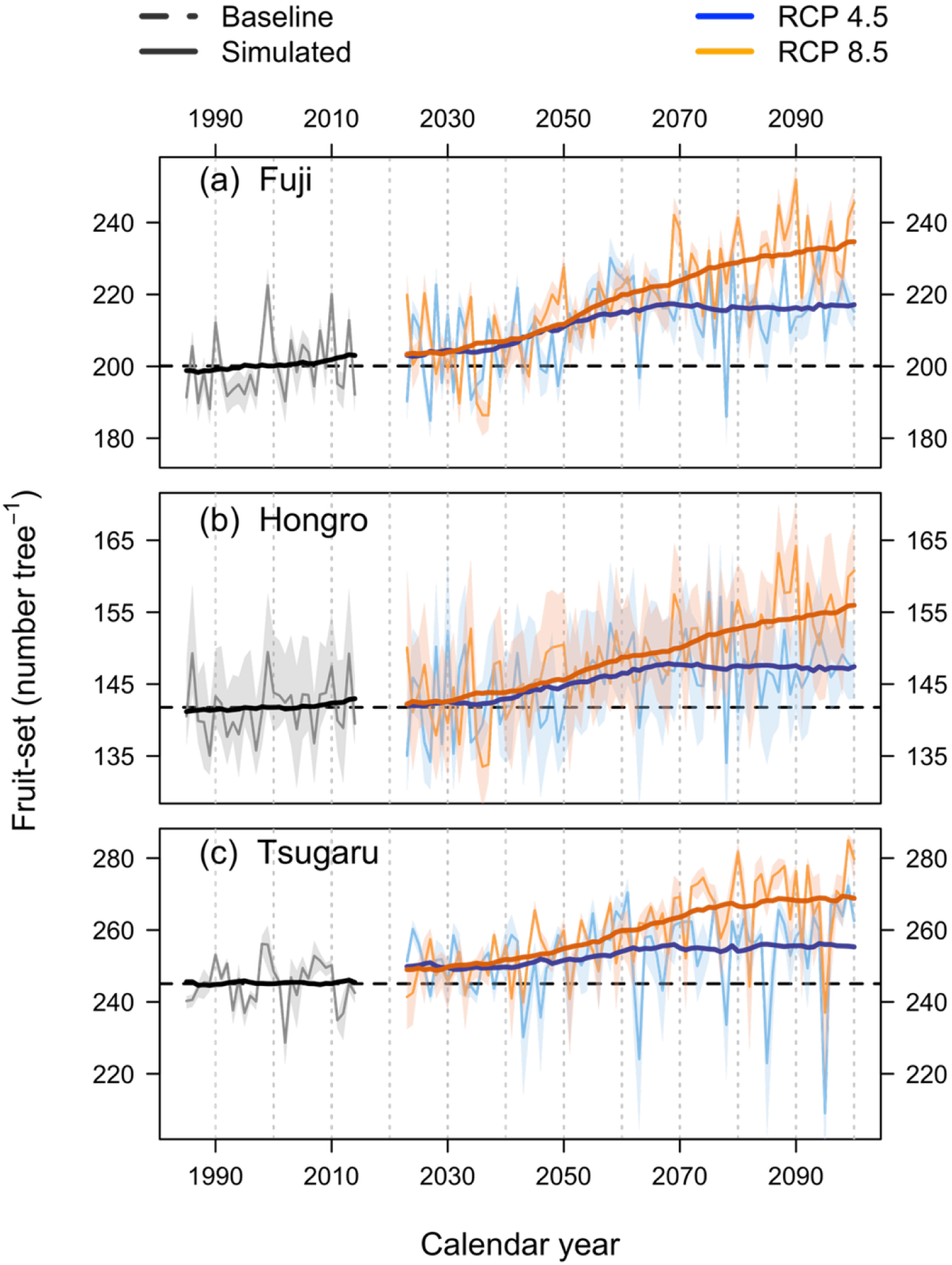
Predicted trends in annual per capita fruit-set under two different climate change scenarios for the period 2023 to 2100. Projected values of future climate were passed to the best-supported models for each cultivar to estimate changes in fruit-set for individual trees in surveyed Korean orchards. Predicted individual-tree yields were then averaged across all orchards for each cultivar and plotted as annual-scale mean fruit-set (colored narrow lines). Temporal trends in future fruit-set were derived from 30-year moving averages (smoothed bold lines). Annual production for a recent 30-year reference period (1984-2014) was also estimated by fitting models with interpolated climate variables (narrow black lines). The bolded black line is the matching trend. Horizontal dashed lines are mean 30-year fruit-set levels for the reference period: 200, 142, and 245 fruit-set number tree^−1^ for Fuji, Hongro and Tsugaru, respectively. Shaded regions are 2-unit support intervals.

Our future simulations predict, in addition to greater overall average productivity, higher levels of interannual variability. Interannual variation in fruit-set (SDΔfruitset, Appendix A9) is estimated to increase marginally in Hongro and more strongly in Tsugaru, particularly under RCP4.5. Production variability in Fuji is comparatively consistent for all simulations. However, individual years of anomalously low production are forecast for all cultivars (Fig. 5). For example, productivity declines exceeding 1.5 SDs below overall mean fruit-set levels are predicted for all genotypes (N=8 years for Fuji and Hongro, and N=6 for Tsugaru).

Our analyses also indicate that temporal changes in future productivity will vary geographically. Spatial variability is more pronounced in Tsugaru (Fig. 6) relative to both Fuji and Hongro (Appendix A10). In southeastern regions, Tsugaru fruit-set levels are predicted to increase twofold relative to reference baseline levels, while in northwestern coastal areas they are forecast to decline by up to 50% relative to baseline conditions.

**Figure 6.**
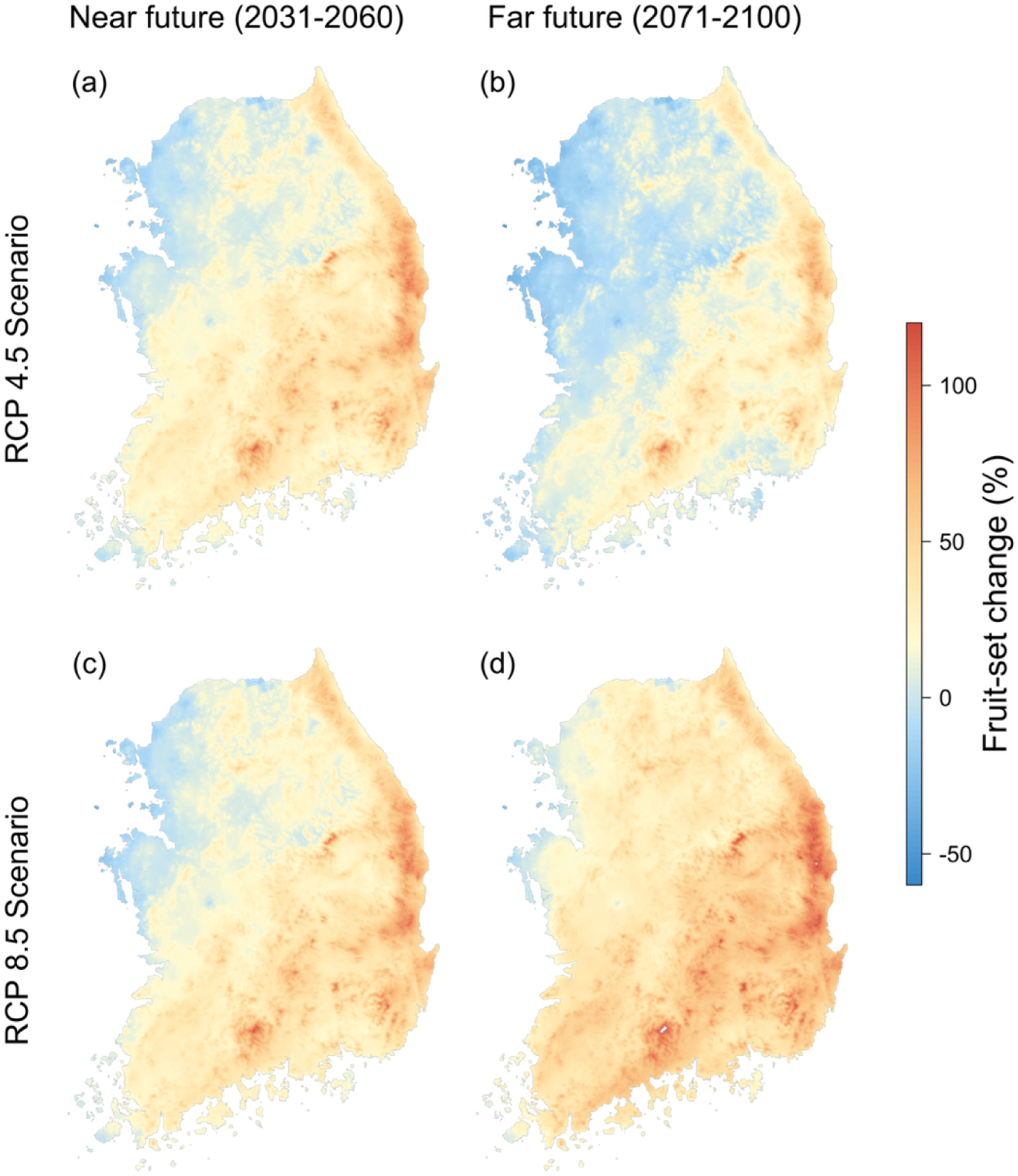
Geographic variation in predicted average annual per capita fruit-set for Tsugaru as a consequence of projected future climate conditions. Data from two climate change scenarios were used in the simulations: an intermediate (RCP4.5) and a high emission (RCP8.5) scenario. Spatially-explicit changes in 30-year average fruit-set – estimated for 30-m pixels based on associated seasonal climate conditions and then averaged for a given 30-year period – for two future periods are shown relative to a recent 30-year base period (1984-2014). Differences in predicted fruit-set (%) between future and reference periods were plotted on a pixel by pixel basis. Tree age and management effects were held constant.

## Discussion

### Fruit-set dynamics

In general, our analyses are consistent with the long-recognized phenomenon of temporally fluctuating fruit production in farm crops, commonly referred to as biennial or alternate bearing (Jonkers, 1979; Monselise and Goldschmidt, 1982; Goldschmidt and Sadka, 2021). The first-differenced chronologies of fruit-set exhibited moderate levels of negative serial autocorrelation (-0.36 to -0.38, Table 1), diagnostic of a reversal of reproductive effort over successive years (Koenig and Knops, 2000). The magnitude of annual fluctuations was modest, not exceeding 15% in most years relative to the overall mean level for a given cultivar and orchard group (Fig. 2). However, we did not observe unambiguous evidence that production followed a strictly bimodal cycle, as has sometimes been observed in prior work (e.g. Sharma et al., 2019; Zakalik et al., 2024). The frequency distributions of annual fruit-set in this study departed from normality by Anderson-Darling tests (p < 0.001) and were marginally right-skewed, but nevertheless clearly unimodal. In summary, our findings suggest that production patterns were not precisely regular, periodic, or bimodal, but instead oscillated around mean levels.

Reproductive effort that is variable across years appears to be an intrinsic attribute of the life-history strategies of many plant taxa. A large literature has examined the dynamics of fruiting in species of natural environments and uncovered ubiquitous, though not universal, evidence for alternate bearing (Koenig, 2021). The coefficient of variation (CV) has been widely used to quantify the properties of seed production in those studies (Silvertown, 1980; Kelly, 1994). A prior meta-analysis of seed production documented an overall mean CV of ∼1.253 for forest trees, based on 474 records for multiple taxa (Kelly and Sork, 2002). In agricultural systems, the innate tendency for variable reproductive behavior across years may be expected to be dampened, via human subsidies, artificial selection, and other cultivation practices (Garcia et al., 2021). Accordingly, CVs of fruit set for apple in Korean orchards are markedly lower relative to values for forest trees, ranging from 0.25-0.34 for orchard groups (Fig. 2). Nevertheless, at local farm scales, maximum CV values approached 0.6, suggesting a persistent tendency for sizeable levels of production variability.

### Mechanisms

The physiological underpinnings of fluctuating reproductive effort in food crops and plants generally have not been clearly resolved (Samach and Smith, 2013; Goldschmidt and Sadka, 2021; Belhassine et al., 2022). A prevailing theory in the ecological literature builds on the high resource costs of reproduction (Bell, 1980; Obeso, 2002). Specifically, constraints to fruiting arise from the simultaneous resource (carbon, water, nutrients) demands of multiple biological processes, including growth, defense, storage, and maintenance metabolism (Herms and Mattson, 1992). Irregular rates of reproduction, despite the fitness costs associated with lost opportunities for progeny recruitment, are assumed to be a necessary response to inherently finite substrate availability. Mathematical models predict that the quantity of fruit-set in a given year in an individual plant is determined by the magnitude of surplus carbon exceeding particular thresholds; thresholds being determined by essential minimum allocation to critical growth and maintenance functions (Isagi et al., 1997; Satake and Iwasa, 2000; Lyles et al., 2015). Fruit production levels are therefore assumed to depend on the resource balance of a plant, which are influenced by varying, climate-modulated rates of accumulation in prior years (Silvertown, 1980), as discussed in the following sections. The resource budget model is consistent with the nutritional resource hypothesis of the horticultural literature (Goldschmidt and Sadka, 2021), which similarly predicts that alternate bearing is fundamentally an outcome of the depletion of photosynthate reserves during years of high fruit output. A complementary theory, though controversial (Guitton et al., 2012; Goldschmidt and Sadka, 2021), involves hormonal controls on reproductive effort. Specifically, gibberellins exported from the seeds of developing fruits, perhaps functioning in concert with auxin, are posited to inhibit the concomitant induction of floral buds, thereby limiting potential fruit-set in the subsequent year (Tromp, 1982; Mutasa-Göttgens and Hedden, 2009). Gibberellin concentrations that increase with crop loads may therefore contribute to alternating production.

### Reproductive synchrony

We identified evidence for moderate levels of spatial correlation in the temporal fluctuations of fruit-set among disjunct orchards. Demographic rates that are synchronized among populations is a prevailing phenomenon in biological systems (Liebhold et al., 2004). Though evidence that reproductive output is cyclical *and* correlated among individual plants of distant populations is ubiquitous in natural environments (Koenig and Knops, 2000), relatively less is known about the prevalence, scale, or consequences of correlated production in agricultural settings (Pearse et al., 2020). Garcia et al. (2021), in a global analysis, identified a persistent signal of alternate bearing for multiple crops in published national-scale chronologies of yield, which they argued was evidence of synchronized fruiting patterns among farms at regional scales. In this study, significant levels of synchrony were detected over distances of up to 25 km, depending on cultivar (Appendix A5). The degree of synchronization was weakest for Tsugaru, which had the smallest dataset in terms of surveyed orchards, yet was nevertheless significant within a range of ∼15 km. We posit that synchrony detection depends in part on sample size, as well as the intensity of sampling within landscapes. We suggest, therefore, that our analyses may underestimate the actual magnitude of synchrony at local scales, and/or the extent of synchrony at landscape scales, as our observational datasets were derived from a stratified random selection of orchards, rather than a comprehensive survey of all farms.

Often referred to as a masting phenomenon, a long history of ecological research has been devoted to deciphering the drivers of reproductive output that is both cyclical and synchronized (Crone and Rapp, 2014; Koenig, 2021). In forests, evolutionary processes are commonly hypothesized; for example, that synchronous flowering among trees may confer a selective advantage due to more efficient density-dependent pollination, or that intermittent but contemporaneous fruiting enhances the probability of either seed dispersal or survival (via escape from seed predators) during large crop years (Janzen, 1971; Smith et al., 1990; Kelly, 1994). Presumably, natural selection factors have been largely attenuated in intensively-managed crop systems. However, variation in the environment, through a so-called Moran effect, has been attributed to correlated metabolic production in plants of both natural (Koenig and Knops, 2013) and agricultural (Rosenstock et al., 2011) systems. More specifically, exposure to extreme or anomalous weather (e.g., drought, frost, warm winters) is proposed as a mechanism that can align reproductive output in trees (Pearse et al., 2016; Garcia et al., 2021, 2024). Extreme events, often termed vetoes or shocks, may inhibit flowering or fruit-set in many individuals in a given population or orchard, even if, by the resource budget hypothesis (discussed previously), the corresponding carbon stores of plants exceed minimum thresholds needed for allocation to reproduction. If such a shock occurs during a year of high fruiting potential, the outcome may be an alignment in future years of the reproductive schedules of plants that were previously out of phase (Lyles et al., 2015; Garcia et al., 2024). As climate is generally correlated over large areas, extreme events have the potential to synchronize distant farms. Consequently, farm-level fluctuations in production may ripple out to affect and potentially destabilize the supply chains of agricultural products at regional scales.

### Climate relations

Models provide evidence that, controlling for other factors, temperature increases during cold-season months may lead to reductions in fruit-set potential. These results are consistent with extensive evidence in the literature that warming disrupts the regulatory effects of chilling and forcing on dormancy progression, with consequences for patterns of budbreak, flowering dynamics, pollination success, and ultimately the magnitude of fruit-set (Inouye, 2022). Observations of phenological advances (earlier budbreak relative to historical patterns) are common and are assumed to be caused by enhanced forcing (Legave et al., 2013; Drepper et al., 2020; Cho et al., 2021; Lee et al., 2023). However, delays in flowering have also been documented and attributed to insufficient chilling (Wang et al., 2020; Vanalli et al., 2021; Medda et al., 2022; Chen et al., 2025). Warming-induced shifts in dormancy release, whether accelerated or impeded, are known to have negative consequences for production (Goeckeritz et al., 2023; Petri et al., 2024; Roussos, 2024; González-Martínez et al., 2026). In the event of advancement, greater frost exposure can cause floral abscission and a limitation of fruit-set (Chen et al., 2025). A protracted dormancy period can delay spring leaf development and associated photosynthetic activity, thereby limiting carbohydrates available for flowering and fruiting (Hoch et al., 2013; Lordan et al., 2019; Fraga et al., 2021). More generally, a phenological shift in either direction may extend the period of anthesis and desynchronize flowering among trees, leading to inefficient pollination (Luedeling, 2012; Ramírez and Davenport, 2013; Tominaga et al., 2022; Fernandez et al., 2023). Temporal niche shifts may further compromise fruit-set through a decoupling of the period of anthesis in trees from the life cycles of wild insects (Peñuelas et al., 2002; Kőrösi et al., 2018), which provide significant pollination services that augment or even exceed the pollen-delivery efficiency of managed honeybees (*Apis mellifera* L.) (Garibaldi et al., 2013; Olhnuud et al., 2022; Sáez et al., 2022; Eeraerts et al., 2025).

The objective of this study was not to model dormancy release, nor the influence of climate on chilling or forcing processes *per se*, the relative importance of which constitutes an ongoing debate (Ettinger et al., 2020; Wang et al., 2022). An explicit modeling of dormancy normally involves the formulation of cultivar-specific, empirically-based chill and heat sub-models that depend on theoretical or experimentally-derived temperature thresholds (Fadón et al., 2020). However, climate factors that determine chilling and forcing conditions are often statistically correlated in space (Ettinger et al., 2020). An increasing body of evidence also suggests that these processes may not only overlap in time but interact functionally so that, for example, forcing requirements may vary as a function of prior chilling states (Basler and Körner, 2014; Pope et al., 2014; Chuine et al., 2016; Luedeling et al., 2021; Delgado et al., 2025). Consequently, we did not attempt to quantify or differentiate these factors in our analyses of fruit-set. Instead, we fit models with *a priori* selected temperature variables corresponding to the coldest months of the dormancy period. We do argue that a parsimonious interpretation of our results is consistent with widespread expectations that winter warming may obstruct or alter dormancy progression (e.g., by diminishing chilling effects) and thereby depress fruiting potential (Atkinson et al., 2013; Inouye, 2022; Meza et al., 2023). Although, photoperiod is expected to play a role in regulating dormancy, some taxa including apple have been found to be insensitive to daylength variation (Heide and Prestrud, 2005). The evidence for divergent, cultivar-specific fruit-set responses identified by our analyses does further, albeit indirectly, support a hypothesis for the importance of chilling. Specifically, the unique positive fruit-set response in Tsugaru to cold-season warming can be assumed to be underlain by the comparatively abbreviated chilling requirements of this cultivar relative to Fuji and Hongro (Pertille et al., 2021; Petri et al., 2024).

Research suggests that in the final phases of dormancy bud development and associated metabolic costs rely on the translocation of carbon from storage organs (Loescher et al., 1990; Basler and Körner, 2014; Fadón et al., 2020). We therefore hypothesized that conditions promoting photosynthesis and reserve accumulation in prior seasons may support a greater fruit-set potential, as has been suggested for forest trees (Buechling et al., 2016). This premise is consistent with the resource budget hypothesis (Isagi et al., 1997) discussed above; specifically that reproductive processes depend on surplus reserves that are available after the carbon requirements of growth and survival functions have been met. Accordingly, for all cultivars in our study, warmer-than-average conditions during bud initiation year were associated with markedly higher levels of fruit-set in the following spring (Fig. 4). Elevated temperatures can promote net carbon gain in trees by extending the growing season (Carroll et al., 2021) and/or accelerating enzyme-mediated biochemical processes that regulate photosynthetic performance (Saxe et al., 2001). However, warming beyond an optimum may cause protein disfunction and thereby depress assimilation rates (Roussos, 2024), though we did not observe evidence for a threshold response in this study (Fig. 3a). We note that the upper temperature limits (95^th^ percentiles, Table 1) observed during the model-fitting period (2007-2014) were generally below the maximum temperatures simulated for future decades, at least under RCP8.5 (Appendix A2). Therefore, possibly more extreme or novel future conditions may have a potential to exceed the physiological optima of these genotypes, thereby constraining carbon uptake potential.

Our models do capture a unimodal-shaped fruit-set response to precipitation (Fig. 3b). An interpretation similarly based on carbon dynamics would suggest that water limitation may compromise the functioning of both xylem- and phloem-based vascular systems. Water stress can cause progressive losses in xylem transport capacity via tension-induced cavitation, which may then, through stomatal closure, lead to declines in transpiration (Korolyova et al., 2022; Goke and Martin 2022, 2023). Transpiration reductions limit CO_2_ absorption and supply to chloroplasts, with attendant effects on photosynthesis (Wiley, 2020). Otherwise, low turgor pressure may limit the translocation of assimilated sugars through phloem from source (leaves) to sink organs (buds), directly inhibiting cell division and expansion (Stanfield and Bartlett, 2022; Novick et al., 2024). Thermal conditions may also interact with water supply to affect hydraulic conductance in vascular tissue. Excess moisture can diminish photosynthesis via decreases in irradiance associated with more prevalent cloud cover (Jackson and Palmer, 1977; Samach and Smith, 2013), the effects of which, in terms of fruit-set reductions, can be compounded when accompanied by cool temperatures (Fig. 4). Otherwise, cool temperatures increase water viscosity, which inhibits solvent flow and therefore carbohydrate transport to developing buds (Roderick and Berry, 2001). Our results show that a combination of cold temperatures and low precipitation minimized fruit-set potential (Fig. 4).

The putative effects of prior resource accumulation on fruit-set variability did not, however, extend beyond the year of bud initiation. Specifically, climate two years prior to fruit-set did not appreciably influence production (Fig. 3d). We assumed that conditions in autumn would be differentially important for reserve accumulation, as carbon demand and partitioning to competing plant functions (growth) are reduced in this period (Priestley, 1964). Although both temperature and precipitation effects were tested, only rainfall was retained in the best-supported models. However, the estimated relationships were essentially flat, indicating that moisture effects were negligible. These results are not fully consistent with the resource budget model, but are partly in agreement with a hypothesis suggesting that carbon fixed by current-year photosynthesis may generally be sufficient to support the construction of reproductive structures (Hoch et al., 2013; Breen et al., 2020).

### Climate change outcomes

In general terms, our simulation experiments forecast an enhancement of mean fruit-set capacity at a nation-wide scale (Fig. 5). Important determinants of these patterns are thermal conditions associated with two key stages of the reproductive cycle; namely, cold-season dormancy and bud initiation. Based on our regression analyses, elevated spring temperatures can facilitate enhanced fruit-set by promoting reserve accumulation, while warmer winters may inhibit production by altering the timing of dormancy release. Climate models forecast strong positive warming trends for both winter and growing-season months, particularly under the more extreme emissions scenario (RCP8.5), for which mean temperatures (Appendix A2) are projected to exceed the 95^th^ percentile limits of observed temperature for the model-fitting time period (Table 1). These steep temperature increases underpin the modeled enhancement of crop performance, with the positive effects of warmth during bud initiation apparently outweighing, or being only partially offset, by the negative effects of reduced cold exposure in winter months.

Simulation results further suggest that future patterns of inter-annual variation in fruit-set will be temporally and spatially variable. For the far future period (2071-2100) and under the intermediate climate change scenario (RCP4.5), simulations predict negative rates of fruit-set growth (Δfruitset; Table A2), or in other words, a decelerating trend in production, for both Fuji and Hongro. An only marginally positive fruit-set increase is forecast for Tsugaru, suggesting a potential stagnation of productivity in this cultivar. For Fuji and Hongro, the deceleration of fruit-set growth trends under RCP4.5 are spatially patchy, but particularly evident in central and southern parts of the country (Appendix A10). For Tsugaru, outright declines in production capacity are forecast for northern portions of the country (Fig. 6). Evidence for decelerating growth trends in some globally important food crops has already been identified in recent studies (Kucharik et al., 2020; Gleiser et al., 2021). A number of causal factors have been proposed, including diminishing returns associated with greater inputs of fertilizers, pesticides and irrigation (Tilman et al., 2002; Aizen et al., 2023), and inherent biological limits to further production gains that have previously been attained via technological advances (Garibaldi et al., 2021). Additionally, biodiversity impoverishment (Dirzo and Raven, 2003), driven in part by agricultural expansion and habitat losses, may be expected to impair pollination services and dependent fruit-set potential (Garibaldi et al., 2013; Garratt et al., 2021). In this study, limits to crop performance in the far future period are underlain by climate change; specifically, projected warming trends in spring are expected to plateau after ∼2070 under RCP4.5 (Appendix A2), presumably limiting reserve accumulation in trees, and thus, dependent fruit-set capacity. A potential for decelerating or stagnating fruit-set gains in coming decades is of major societal concern, given the escalating food demands associated with global population growth (Tilman et al., 2011). Notably however, steeper and more prolonged warming trends forecast under RCP8.5 are predicted to maintain positive trends in fruit-set growth through the simulation period (Fig. 5). We reiterate however, that the novel thermal conditions expected under this more extreme climate scenario have a potential to exceed the physiological optima of apple trees, which is not captured by our modeling framework.

In addition to temperature-mediated effects on the magnitude of fruit-set, our analyses suggest a potential for larger future fluctuations in production. According to simulation results, interannual differences in output (SDΔfruitset, Eqn. 5) will be amplified to differing degrees for all cultivars relative to 30-year baseline levels (Appendix A9). Maximum variability is expected under RCP4.5 in the far future period. Under RCP8.5, more marginal increases in variability are predicted, and only for Hongro and Tsugaru. Beyond concerns related to a deceleration of production, a possible amplification of interannual variability may additionally compromise food security, as both farmer revenues and stable food supplies depend on consistent production across years (Garibaldi et al., 2011; Gleiser et al., 2021). Further, our phenomenological-based models do not account for the negative and potentially synchronizing effects of environmental shocks. Extreme weather, such as drought or frost, can not only cause local crop failures in individual farms in individual years, but also align production among disjunct farms and thereby amplify and synchronize crop fluctuations at regional scales (Garcia et al., 2024). Additional investigation using process-based models is needed to understand the possible consequences of novel climates (i.e., beyond the range of our datasets) and extreme events for the physiological performance of trees, synchrony among orchards, and the stability of regional crop supplies.

## Conclusions

Evidence in this study of a negative effect of cold-season warming on fruit-set is largely consistent with widespread expectations for climate change disruptions of dormancy processes and flowering dynamics (Fernandez et al., 2023; Meza et al., 2023). The distinctively positive response of the single low-chill cultivar, Tsugaru, to increasing winter temperatures also indirectly supports a hypothesis for the relevance of chilling conditions in mediating reproductive cycles. Our results highlight the importance of legacy effects, as climate during the year prior to fruiting tempered or even counteracted the negative influence of warm winters. These relationships are the basis of our forecasts for potential production gains as a consequence of future warming, though such outcomes were also accompanied by heightened interannual fluctuations in fruit-set. A potential for correlated production among clusters of farms, as found in this study, may amplify the negative consequences of volatile, unstable fruiting for food supplies. The presence, scales, and causes of synchrony in crop systems in general have not been well quantified, despite the relevance for food availability, and therefore constitute an important ongoing research area. Overall, developing climate adaptation strategies for agricultural production has become a societal priority. Our modeling framework may complement other statistical approaches, such as climate envelope models (e.g., Meza et al., 2023), to inform adaptation strategies that can, for example, guide a selection of cultivars and growing sites that will maximize productivity under future climates.

## Data availability

Model fitting datasets will be available at Zenodo at a future date.

## Authorship contribution statement (CRediT)

Arne Buechling: Conceptualization, Formal analysis, Investigation, Methodology, Visualization, Writing – original draft, Writing – review & editing. Jung Gun Cho: Conceptualization, Data curation, Formal analysis, Investigation, Methodology, Visualization, Writing – review & editing. Jakob Pavlin: Writing – review & editing. Forzia Ibrahim: Writing – review & editing. Patrick H. Martin: Funding acquisition, Resources, Supervision, Writing – review & editing.

## Acknowledgements

This work was carried out with the support of “Cooperative Research Program for Agriculture Science & Technology Development (Project No. PJ012116)”, Rural Development Administration, Republic of Korea. We also thank NongHyup Property and Casualty Insurance for providing the orchard data and the Korea Meteorological Administration for supplying weather data.

## Supplementary materials

Appendix A1: Range in climate conditions associated with Korean orchards

Appendix A2: Projections of future climate conditions in the Republic of Korea

Appendix A3: Frequency distributions of fruit-set

Appendix A4: Temporal autocorrelation in observed orchard production

Appendix A5: Extent of spatial correlation in fruit-set production among orchards

Appendix A6: Plots of observed and predicted fruit-set from regression models

Appendix A7: Parameter values of the best-supported regression models

Appendix A8: Effects of tree age on fruit-set production

Appendix A9: Summary statistics describing simulated trends in future fruit-set capacity

Appendix A10: Predicted responses in fruit-set production to scenarios of future climate

